# Non-Optical, Label-free Electrical Capacitance Imaging of Microorganisms

**DOI:** 10.1101/2024.09.25.615041

**Authors:** Joseph T. Incandela, Kangping Hu, Pushkaraj Joshi, Jacob K. Rosenstein, Joseph W. Larkin

## Abstract

Many fundamental insights into microbiology have come from imaging, which is typically synonymous with optical techniques. However, the sample preparation needed for many optical microscopy methods such as labeling, fixing, or genetic modification, limits the range of species and environments we can investigate. Here we demonstrate the use of electrical capacitance measurements as a non-optical method for imaging live microbial samples. In electrical capacitance imaging (ECI), samples are positioned in contact with a semiconductor sensor array, and localized capacitance measurements are made across the array. From these measurements, we generate textured images of a variety of microbial colonies. We determine that capacitance is correlated with local sample thickness by comparing ECI data to 3D confocal scans. We further illustrate with ECI that a difference in capacitance signal allows microbial species to be spatially distinguished in co-culture conditions. In order to highlight the versatility of our system, we capture the cross-sectional development of floating pellicle biofilms in a liquid culture at millimeter length scales during weeks-long time-lapse experiments. These novel results establish a new low-cost and portable platform which can be used for spatially and temporally resolved experiments in diverse environments with a wide variety of microbial species.

**IMPORTANCE:** Microbes live in diverse environments, and occupy biological roles across many timescales. Investigating the full scope of microbial activity requires imaging systems appropriate to each context. Though optical microscopy is powerful, the use of light, lenses, and other hardware limits where it can be applied. At the same time, existing non-optical imaging methods are frequently destructive to samples and require extensive equipment. In this paper we present a non-optical imaging system that is small, cheap, requires no sample labeling, and is compatible with a variety of microbial species. Our system uses semiconductor chips to measure the inherent material properties of a sample with spatial sensitivity, producing images of microbes contrasted against their environment and each other. Our technique captures label-free, micrometer-resolution images with a pocket-sized device, enabling microbiological imaging experiments in new environments with new species.

## INTRODUCTION

Imaging methods occupy a central role in the study of microbial communities, and have revealed much about their development. These methods have been critical to the exploration of large-scale community dynamics such as chemotaxis in populations of motile bacteria [1], genotypic segregation in expanding colonies [2, 3], and the structure of bacterial biofilms [4]. Microbiologists are increasingly interested in experimental systems that investigate key features of the natural habitat, including non-model species [5, 6, 7], multi-species communities [8], and lab environments that mimic wild conditions [9, 10, 11]. Fluorescence microscopy is the standard optical imaging approach in all of these systems. It is extremely powerful, but requires compromises that limit the data that can be collected. Labeling methods are often destructive [12, 13] which prevents their use on live samples, or cannot be applied to non-genetically tractable species [14, 15]. In fluorescence imaging techniques that can be applied to live samples, phototoxicity and photobleaching limit the length and temporal resolution of time-lapse experiments [**Isha2017, Diaspro2006**] and the anoxic conditions present in many microbial environments hinder the use of fluorescent proteins [16]. For these reasons, among others, researchers have called for new imaging techniques in microbial systems that can observe multiple species over large length scales in diverse environments [17].

Non-optical imaging approaches have the potential to overcome some limitations of optical microscopy, as they do not require labeling and utilize different hardware. Microbial imaging with electron microscopy [18] or atomic force microscopy [19] can reveal high resolution details, but they are difficult or impossible to apply to live samples. Mass spectrometry imaging can resolve chemical and phentoypic distributions within microbial communities [20], but requires sample destruction and highly specialized equipment. A promising non-optical imaging approach that is less invasive relies on arrays of complementary metal–oxide–semiconductor (CMOS) transistors. CMOS sensor arrays, which have been deployed for a variety of biotechnological goals [21, 22, 23], can perform non-optical Electrical Capacitance Imaging (ECI). In this technique, a sample is positioned in direct contact with a CMOS array and a localized measurement of electrical capacitance is made at each sensing pixel. The measured capacitance varies with the local chemical composition of the sample [24]: cells, extracellular material, air, and aqueous media all have different dielectric properties [25]. This allows CMOS arrays to generate contrasting electrochemical images of biological samples based on their inherent electrical properties. Previous researchers have used capacitance measurements to track cellular growth [26], distinguish healthy and cancerous mammalian tissue [27], and to infer the growth phase of sulfate-reducing bacteria [28]. However the use of ECI to capture dynamics at the scale of a microbial colony requires a microelectrode array capable of generating large field of view images with sufficiently high resolution [29].

In this work we use a 5.12 mm *×* 2.56 mm CMOS array to perform non-optical capacitance imaging of microbial communities. We collect ECI data of samples brought into contact with the CMOS array [29], and compare our electrochemical imaging data to corresponding fluorescent light (FL) images. We demonstrate an ability to generate high contrast images that resolve spatial features of microbial biofilms, by exploiting their distinct and spatially variant capacitance signal. We show our method can be used to study diverse species and note that each species and its preferred growth media exhibit a distinct range of capacitance values. To investigate the relationship between sample capacitance and geometry, we compare the distribution of capacitance in ECI data to the biofilm thickness distribution determined with confocal z-stack imaging. Furthermore, with ECI we demonstrate the ability to distinguish multiple microbial species in the same field of view, and compare to multichannel fluorescence data. Finally, we showcase the flexibility of this system and it’s ability to integrate with diverse culture formats, by imaging the profile of pellicle biofilm development at liquid-air interfaces during weeks-long time-lapse experiments. Our method occupies a new niche in microbial imaging: it is compact, cheap and label-free with a high frame rate and large field-of-view. With this technique, new microbiological imaging experiments will be possible. For example, our ECI can readily be used for non-model or untractable species because it forms images based on innate sample material properties and requires no labeling; the physical robustness of our sensor will enable weeks-long time lapse imaging experiments, and the low cost and small size of our sensor will facilitate its use for imaging in the field.

## RESULTS

### High-contrast capacitance imaging reveals biofilm structure

Our imaging system consists of a CMOS chip mounted on a 4 cm *×* 4 cm circuit board (Fig. 1A, left), which in turn is connected to a data acquisition board (Fig S1). The data acqusition assembly provides power and control signals to the sensor chip, and connects to a computer through an FPGA module and USB interface [29]. The CMOS array itself contains 131,072 sensing pixels arranged in a 512 *×* 256 rectangular grid, which results in a 13.1mm^2^ field of view during imaging (Fig 1A, center). Finally, each of the pixel sensors consists of a 10*µ*m *×* 10*µ*m titanium nitride electrode (Fig. 1A, right), giving a spatial resolution of 10*µ*m, which can potentially be improved algorithmically [30]. While biomass is positioned in contact with the CMOS surface (Fig. 1B), circuits within the chip can measure the local capacitance at each pixel (Fig. 1C) by applying AC voltage waveforms and measuring the capacitive charge response [29]. Inclusion of an external bulk reference electrode can be helpful in some configurations, but the AC-coupled ECI measurements can be performed without a reference electrode (Fig. S4) [29, 31]. The capacitance measured at each pixel depends on the geometry of the conductive electrode and the dielectric properties of the material near it. The geometric contribution is nominally the same across the array pixels, yielding capacitance values that vary predominantly with the local material properties of the sample.

**Figure 1.**
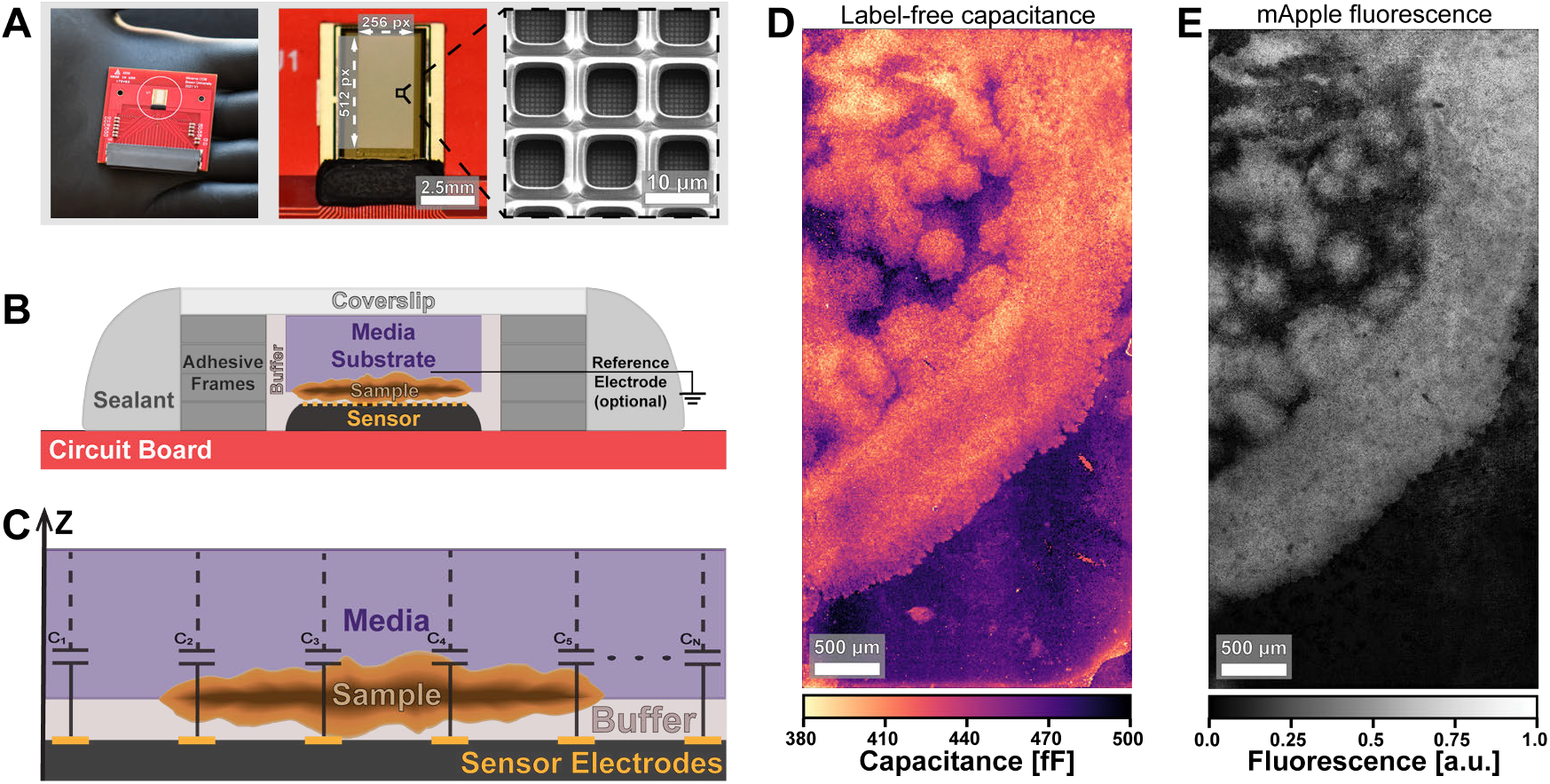
A compact, non-optical imaging system for microbial samples. (A) A photo of the CMOS chip held in hand (left), a closeup of the sensor array (center), and an electron microscope image of the sensing pixels (right). (B) A schematic of the sample setup and (C) the spatial capacitance measurement. (D) A capacitance image of a *B. subtilis* biofilm (strain 3610Δ*sinR*) and (E) a fluorescence image of the same sample. The fluorescent label is a constitutively expressed mApple fluorescent protein.

To demonstrate our system’s ability to take images of microbial samples, we mounted a *Bacillus subtilis* biofilm (matrix overproducing strain NCIB3610Δ*sinR*, which creates colonies with notable spatial structure, Table S1) on the sensor and took both capacitance (Fig. 1D) and fluorescence images (Fig. 1E).

The biofilm’s textured features can be observed with either modality, however there are notable differences in the images. For example, ECI reveals colony edges in finer detail. It is important to note that ECI and FL capture images from opposing sides of the sample (Fig 1B). Biofilm samples were therefore prepared to a thickness *≤* 40*µ*m to allow fluorescence to be collected from the full sample thickness. However, we anticipate important differences between the datasets that are tied to the nature of these imaging modalities. One fact that accounts for differences in capacitance and fluorescence images is that fluorescence imaging detects signal from fluorescent proteins in cells, while ECI measures capacitance from all biomass, including cells and extracellular matrix. Fluorescence in Fig. 1E comes from the constitutive expression of a fluorescent protein. Signal intensity in the image thus depends on cell density and protein expression, both of which may vary across the colony [**Gaigalas2001**]. However in the electrical capacitance measurement of Fig. 1D, the cells and extracellular biomass both have distinct electrochemical signatures, which are contrasted against the background of higher capacitance culture media [24].

### Imaging diverse species with ECI

ECI relies on inherent material properties and can image any biological sample as long as the dielectric properties provide sufficient contrast. The method can thus be used to image wild-type species with no genetic modification, in their preferred growth media. In Fig. 2, we present measurements of the undomesticated *Bacillus subtilis* strain PS-216 [**Durett2013**], a biofilm-forming strain of *Bacillus licheniformis* [**Jespersen2012**], as well as a strain of *Saccharomyces cerevisiae* (photos of sample colonies in Fig. 2A; more strain information in Table S1). In Fig. 2B and Fig. 2C we compare fluorescence (Thioflavin T for *B. subtilis* and *B. licheniformis*, mNeonGreen for *S. cerevisiae*) and capacitance images of these samples, illustrating the high-contrast, non-optical images attainable for any many microbial samples. The images from each modality have some differences, for example there are regions where we observe colony signal in ECI but not in fluorescence. Again, we hypothesize that these differences are from fluorescence imaging detecting signal primarily from labeled cells, while ECI is detecting signal from all biomass, including cells and extracellular matrix.

**Figure 2.**
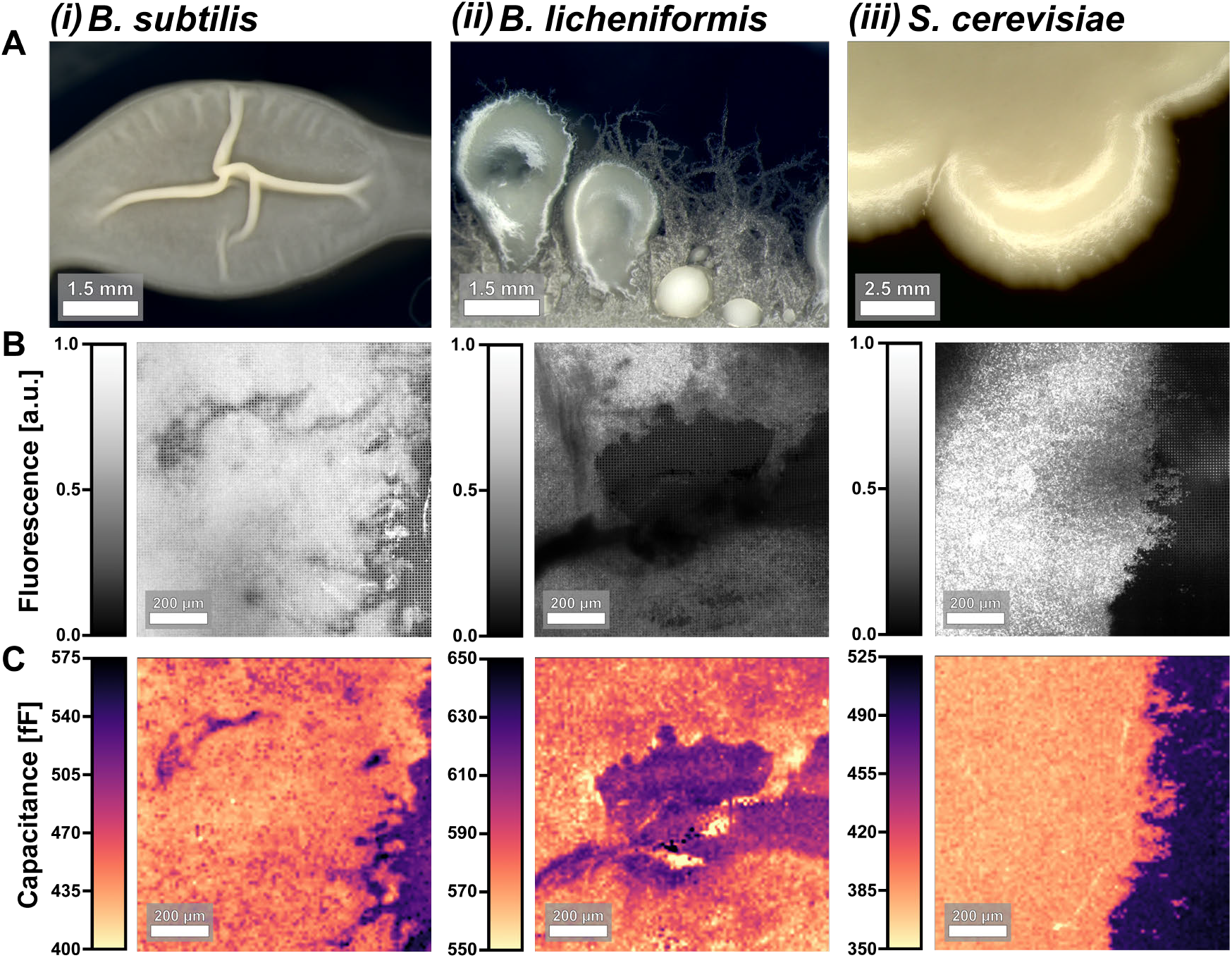
Capacitance imaging of multiple microbial species. (A) Photographs of colonies grown on agar substrates. (B) Fluorescence and (C) capacitance images from the same field of view in a sample of each species. Note, these are not from the same samples photographed in (A). The species represented are; (i) *B. subtilis*, (ii) *B. licheniformis*, and (iii) *S. cerevisiae*. Fluorescent reporters are Thioflavin T for (i) and (ii) and mNeonGreen for (iii).

Unlike fluorescence microscopy, ECI measures a quantity with physical units, namely capacitance. For this reason, the substrate becomes part of the image signal in a different way from fluorescence imaging. We highlight this by noting that the ECI data in Fig 2C contains distinct capacitance ranges due to the images having different species, but also different media substrates (Table S2). It is worth noting that the range of absolute capacitance values (Fig. 2C) can vary across species and growth media (Table S2). Biomass tends to have lower absolute capacitance than the surrounding culture media, independent of the species/media combination.

### Capacitance correlates with sample thickness

We anticipated that capacitance values measured with ECI would depend on the local thickness of the sample. At the same time, we expected that screening from ions in the media would set an upper limit on ECI’s sample depth sensitivity. To test our ability to interrogate 3D sample geometry, we compared ECI data of a *B. subtilis* biofilm (Fig. 3A), to a thickness map of the biofilm (Fig. 3B) extracted from a 3D confocal scan (Fig S5). Capacitance is negatively correlated with biofilm thickness, as seen in the line profile of Fig. 3C. We expect a negative correlation because bacterial biomass has a lower permittivity than aqueous media [32]; thick regions of the biofilm have low capacitance and thin regions have higher capacitance.

**Figure 3.**
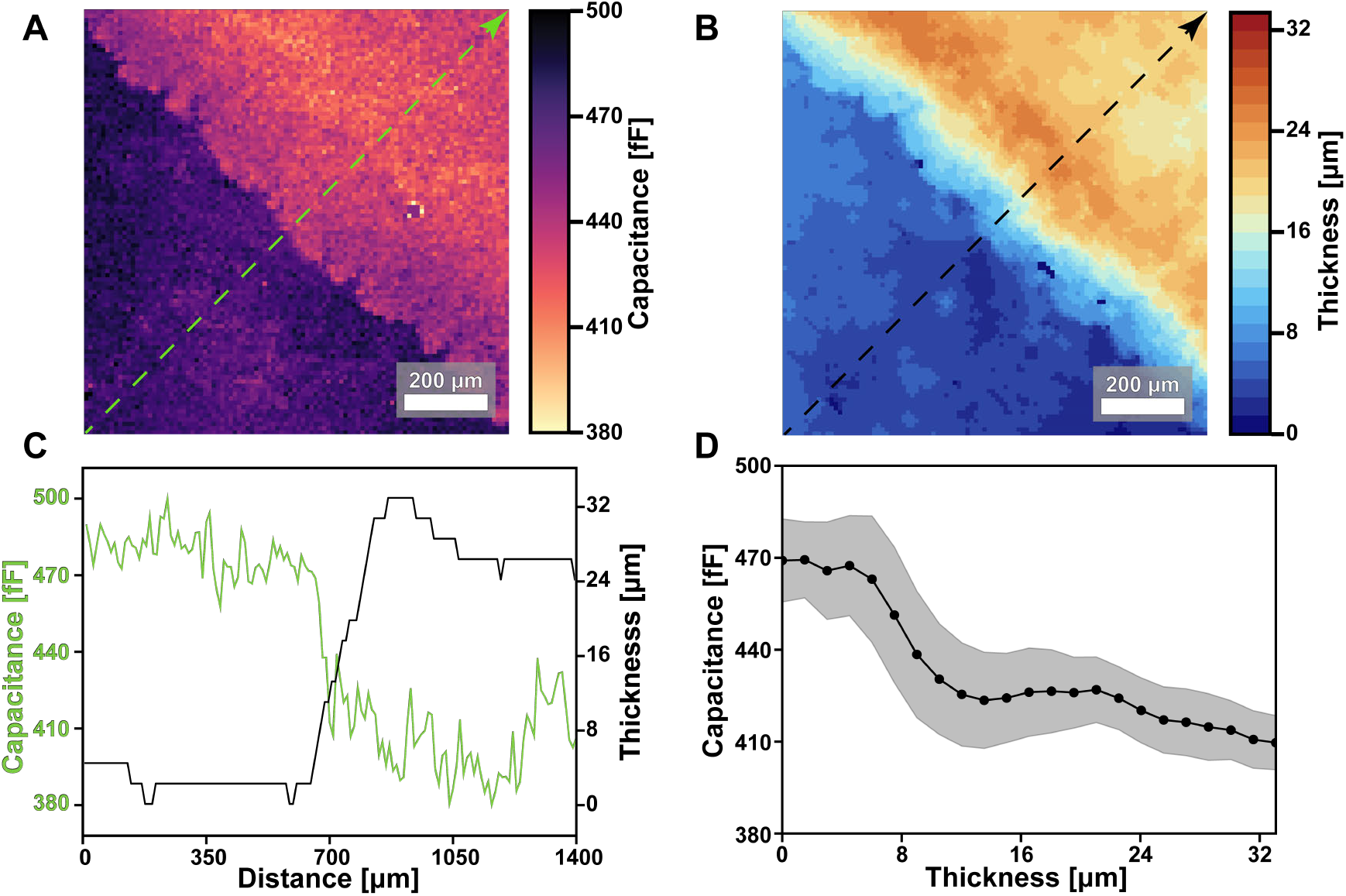
Relationship of capacitance to sample thickness. (A) A capacitance image of a *B. subtilis* biofilm edge and (B) the corresponding thickness map determined from 3D confocal imaging of mApple fluorescence. (C) Line traces of sample capacitance (green) and thickness (black) along the line profiles of (A) and (B). (D) Capacitance values taken from each position across the full 13.1 *mm*^2^ area of the CMOS array are plotted against the corresponding local thickness.

To determine the sensitivity to sample thickness, we performed a pixel-wise comparison between every point in the 512 *×* 256 capacitance image and the corresponding value in the thickness heatmap. From the segmented confocal data, we measured biofilm thicknesses ranging from 0 to 33.2 *µm* across the sample. The measured thicknesses were discretized into 22 values due to our confocal z step size (Fig. S5). In Fig. 3D we plot the mean and standard deviation of the capacitance values matching each of the 22 z positions. We observe a non-linear correlation between capacitance and thickness with an apparent plateau for sample thicknesses ≥ 16 *µm*. This analysis suggests that while our ECI measurements could continue to detect biomass thicker than 16 *µm*, beyond that point the capacitance values may not change significantly with sample thickness.

This analysis helps establish our ability to measure local thickness, and reveals its limitations. However, the ECI data presented here is measured in a self-capacitance configuration, which is not the only configuration available to our sensor array (Fig. S4). Similar correlations between sample geometry and mutual-capacitance datasets are possible, and have been used to improve machine-learning approaches for computing 3D biomass distributions from spatial capacitance measurements [31].

### Mixed species capacitance imaging

We wanted to know if we could use our method to image the colonies of multiple species within the same sample. as capacitance signal depends on sample geometry and the inherent dielectric properties of a colony. Based on our single-species imaging data, we thought that differences in species dielectric properties or colony geometry could be sufficient to spatially segment species within a multi-species sample using ECI. To explore this possibility, we inoculated the same substrate with *B. subtilis* expressing *mCherry* and *S. cerevisiae* expressing *mNeonGreen*, and took both ECI and FL images (see Materials and Methods for strains and conditions). In the ECI data, it is apparent that there are two regions with distinct morphologies (Fig. 4A). The boundary between them is clear even on small scales (Fig 4B). From the corresponding multichannel FL data (Fig 4C), we confirmed the boundary between the two species and identified the species in each population (Fig. S6). By applying a species mask derived from the patterns in the ECI data (Fig 4A), we segmented the ECI image into regions classified as *B. subtilis* or *S. cerevisiae* (Fig. S6). We plot the histograms of capacitance measurements within those two regions as well as the whole chip field of view in Fig. 4D. The capacitance distributions from each region are cearly distinct, suggesting that the species are not only morphologically distinct under these conditions, but that they can potentially be separated by their characteristic capacitance ranges.

**Figure 4.**
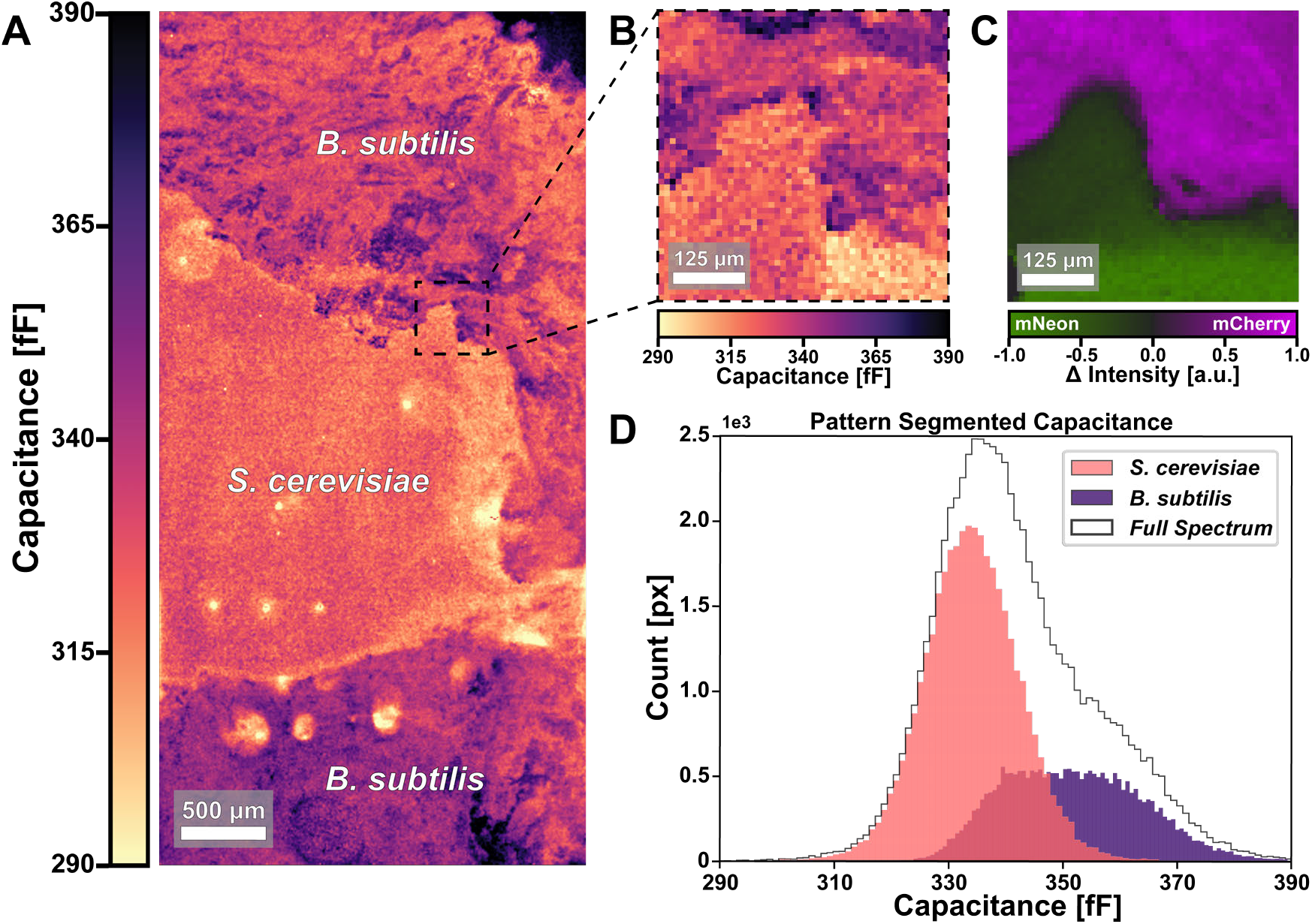
Imaging multispecies colonies with ECI. (A) An ECI image of a sample with *S. cerevisiae* and *B. subtilis* exhibits clear boundaries between their populations. (B) A 1 *mm*^2^ cropped region at the apparent boundary and (C) the corresponding multichannel fluorescence data validating the population boundary. *S. cerevisiae* expresses *mNeonGreen, B. subtilis* expresses *mCherry*. (D) A histogram of capacitance values shows distinct distributions for the two populations, when segmented using the morphological boundaries apparent in (A).

### In-situ ECI Enables Long-Term Monitoring of Pellicle Formation

The small size of our imaging chip facilitates its incorporation into many microbial growth formats. To exploit that capability, we created a system in which the CMOS chip forms one side of a liquid culture tube to image cross-sections of developing colonies (Fig. 5A). When we inoculate the liquid media with *B. subtilis* cells, they form a floating pellicle biofilm at the liquid-air interface [33]. Due to the thickness and opacity of pellicles, researchers have typically resorted to custom microscopy systems to image them [34, 35]. In our system, the imaging chip is directly integrated into the pellicle growth chamber. We set up the sample such that the liquid-air interface would be just above the sensing surface (Fig. 5A), but our sensor can be used to image this interface as well (Fig. S8). By taking capacitance scans at regular time intervals, we were able to image pellicle cross sections as biofilms grew and developed over many days (Fig. 5B). The distinct contrast between biomass and liquid media allowed us to track the area of the sensor colonized by the pellicle over time (Fig. 5C). Within the colonized region of the sensor, we were able track morphology as the pellicle developed (Fig. 5B,D,S7). The initial suspension of cells transitions to form a pellicle beneath the liquid-air interface, where by hour 30 the pellicle is clearly distinguishable from the surrounding media. Notably, the area of the pellicle remains constant for some time (Fig. 5C), before entering a steady growth regime. We hypothesize that early growth contributes predominantly to increasing the density of the pellicle. This hypothesis is supported by the observation that mean capacitance signal in the pellicle region decreases over time (Fig. 5B,S7C) as we would expect from an increase in biomass. Furthermore, while initially the capacitance signal within the pellicle appears spatially uniform, as the colony area expands we observe distinct horizontal patterns within the capacitance images (Fig. 5B). We quantify the development of these patterns by plotting the variance of measured capacitance within both the pellicle and the liquid media over time in Fig. 5D. The new growth at pellicle’s leading edge exhibits spatial regions of varied capacitance (Fig. S7B). Thus as the pellicle expands, the variation in capacitance signal increases as the biofilm develops regions with more heterogeneous capacitance signal. In contrast, the variance of measured capacitance within the liquid media remains constant throughout the experiment (Fig. S7D).

**Figure 5.**
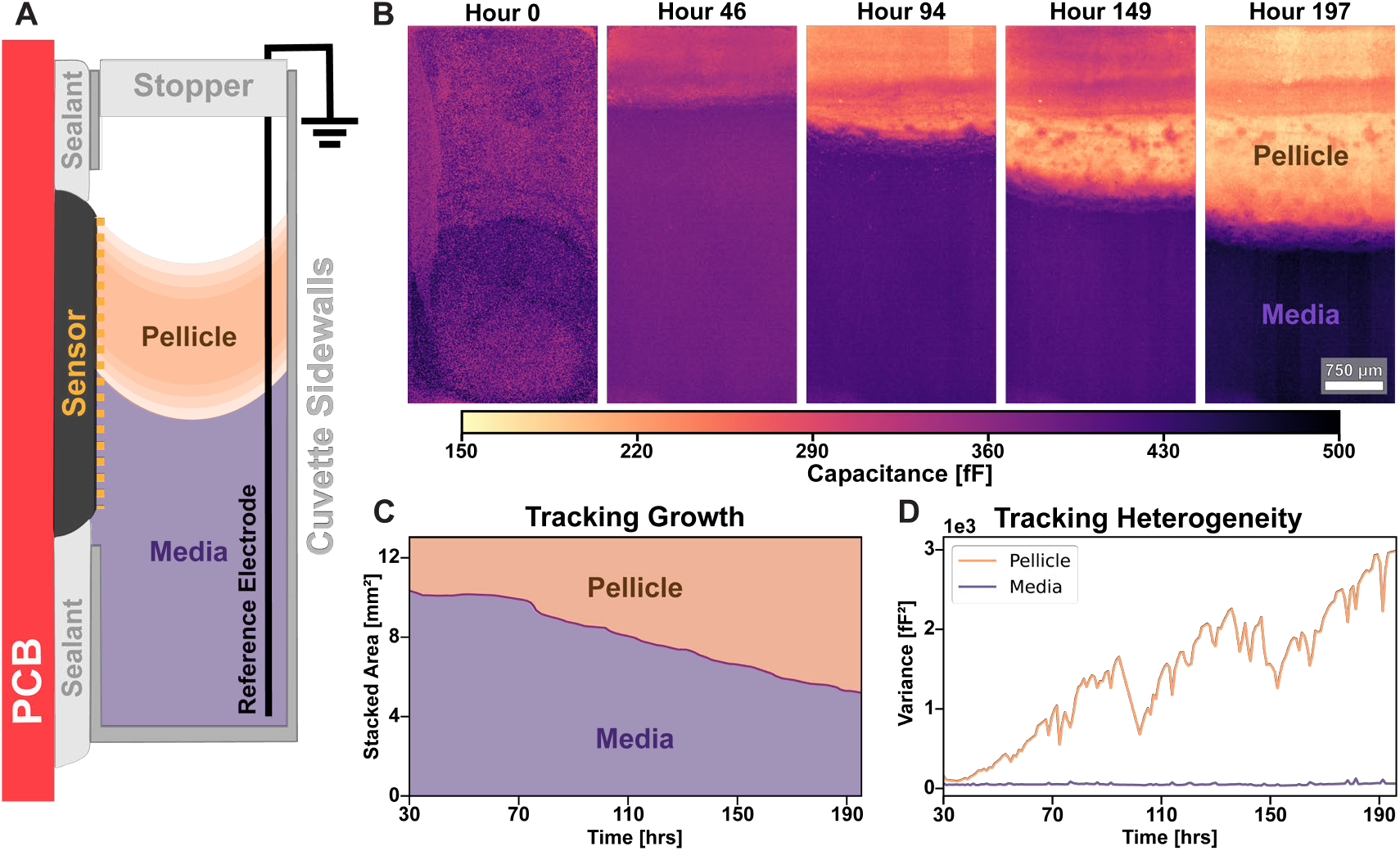
Imaging the profile of pellicle formation for a week. (A) A schematic of the pellicle culturing setup. (B) Capacitance images of pellicle development throughout the time-lapse. Images were taken every 10 minutes, but averaged within 1-hour windows. (C) A stacked area plot of the CMOS array field of view, showing the change in area of the pellicle and media regions. (D) A plot of the capacitance signal variance within the pellicle and media regions over time during pellicle growth. As the pellicle grows and establishes spatially varying capacitance signal, its variance increases, while the variance within the media region remains low and constant throughout the experiment.

## DISCUSSION

We have shown the ability to take images of microbial communities with a handheld device, spanning mm length scales at 10 *µm* resolution, without light. Our system can image a variety of species with no need for labeling. The distinct spatial capacitance signatures of different strains and species enable us to spatially distinguish genotypes in multispecies samples. Our device’s measurements are spatially sensitive to sample thickness up to a cutoff due to ionic screening. Lastly, the compact size of our device enables its integration into a variety of microbial growth systems. We demonstrate this by imaging the cross-section of a pellicle biofilm over the course of several days at 10 minute intervals. We take non-optical images with richer spatial information than previous semiconductor sensors due to the high pixel density of our sensor [36, 37].

Our results demonstrate the proof-of-principle for large-scale CMOS imaging of microbial samples. We anticipate that the system will be used for a variety of new applications that are difficult with conventional imaging. The lack of optical hardware will facilitate *in situ* imaging of environments such as soils, where optics are impractical and ineffective. The ability to spatially map capacitance enables the imaging not only of microbial biomass, but also of mineral substrates [29]. Merging microbial and abiotic solid imaging could enable new microbe-mineral interaction experiments such as observing bacterial growth on specific mineral substrates [38] and monitoring the degradation of tooth substrates by dental biofilms [39]. Combining capacitance imaging with machine learning could facilitate the 3D mapping of species in polymicrobial communities [31]. Exploiting the electrochemical activity of our device’s electrodes [40] could enable simultaneous imaging of biofilms and chemical gradients.

## MATERIALS AND METHODS

### Biofilm sample preparation for ECI and confocal imaging

In the experiments of Fig (1,2,3,4,S4), the biofilms of each species (Table S1) were grown overnight at 30^*o*^C on a 1% agarose substrate of their preferred media (Table S2). For those species without constitutive fluorescence, a 10*µ*M concentration of Thioflavin T (ThT) was added to the substrate for fluorescence. The rest of the sample preparation protocol was designed to meet the challenge of producing biofilm samples that could be imaged first by ECI and later validated via confocal microscopy without any change in the interim.

First, a precise substrate shape was required with an area matching the 2.56mm x 5.12mm CMOS area and with a thickness less than the 2.4mm working distance of the microscope objective. In order to precisely control the substrate shape, two cover-slips were placed on an 85^*o*^C hotplate. At the same time a 1% agarose media mixture was prepared on an 85^*o*^C hotplate with a stir bar to prevent the agarose from setting. *∼*700 *µ*L of liquid agarose media was pipetted onto one of the cover-slips and the second cover-slip was aligned and gently dropped onto the liquid film, causing the still-liquid media to spread between the cover-slips and produce a rectangular slab of agarose media, which could cool and be diced to the desired dimension. This method resulted in sufficiently uniform substrates with a typical thickness of *∼*800*µ*m.

The diced substrate was inoculated with cells grown in a shaken 37^*o*^*C* liquid culture of their respective media. 0.1-0.5*µL* of exponential phase culture was pipetted onto the prepared agarose pad. Inoculated pads were parafilm sealed in a petri dish to reduce moisture loss, and then left in a 30^*o*^*C* incubator overnight. Before loading a sample, the CMOS array was primed with *∼* 200*µ*L of methanol to aide in surface wetting and methanol was carefully exchanged for phosphate-buffered saline (PBS), all without letting the CMOS dry. A mature biofilm sample was then pressed into this layer of PBS, making electrolytic contact with the CMOS. The sample was fixed in place with double-sided adhesive and a coverslip to keep it in contact with the CMOS, and the assembly was sealed with quick-curing silicone (Smooth-On, Inc.) to prevent drying. ECI and confocal imaging was then performed on this completed assembly (Fig 1B).

The use of PBS to prime the CMOS is not necessary for capacitance imaging. Any thin film of electrolytic solution can be used, including the sample growth media. The use of PBS in the protocol permitted an experimental condition favorable to the creation of correlated ECI/confocal measurements, wherein the possibility of changes to the sample’s geometry, electrochemical environment and fluorescent expression was minimized. If measured from the moment a sample was loaded onto the CMOS, the total time required to capture a combined ECI and confocal z-stack dataset was typically 2-3 hours, with confocal imaging representing the bulk of that time. Due to the small volume of the media substrate, cell proliferation in the biofilm was minimal after the overnight growth period, however cells remained viable and expressed fluorescent protein. As a final measure to screen datasets for any changes that may have occurred, ECI and FL data (taken via macroscope) was acquired before and after the long confocal scan.

### Pellicle biofilm sample preparation

For the pellicle growth of Fig 5, *B. subtilis* (NCIB3610) cells were grown aerobically in a shaken 37^*o*^*C* liquid culture of LB into exponential phase (*∼* 1 hour). Cells were then re-suspended in a culture of Minimal Salts glycerol-glutamate (MSgg) and brought back into exponential phase at 37^*o*^C (*∼* 1-2 hours). A 3mL cuvette with a hole cut to introduce the CMOS was adhered to the sensor module with a watertight silicone gasket (Smooth-On, Inc.). The CMOS array was primed by pipetting *∼* 200*µ*L methanol onto its surface, and then rinsed out with *∼* 3 mL DI water several times without letting the CMOS dry, removing the methanol while keeping the CMOS surface primed. Finally, the DI water was exchanged for exponential phase cells in *∼* 2.4mL of MSgg culture, with an OD of 0.2. The volume of media introduced was then reduced to bring the liquid level in the cuvette down to the edge of the CMOS. This allowed growth at the air-liquid interface to begin within the the CMOS sensor’s field of view. To reduce evaporative loss, a cuvette stopper was inserted into the top of the cuvette and subsequently sealed with parafilm. The whole assembly of sensor module and FPGA acquisition board was then placed into an incubator, with the temperature during the experiment typically settling at 32^*o*^*C*. The preparation for the long term pellicle growth featured in Fig. S8 was the identical, however the final assembly was left at room temperature to slow evaporative loss.

### Confocal Validation

All fluorescence microscopy data was taken with a Leica Stellaris 5 Laser Scanning Confocal Microscope, using a LEICA HC FLUOTAR L VISIR 25x/0.95 water immersion objective. An additional 1.28x zoom factor was applied during scans, resulting in 32x total magnification. For each fluorescence dataset, a 512px x 512px image was captured at each position in an 8×15 grid spanning the full CMOS area. The process was repeated at 100 z-positions, ranging from the surface of the CMOS array to 150*µ*m above. A pixel dwell time of 1.2125 *µ*s was achieved with a 600Hz, Bidirectional X scan, and 2x frame averaging. A 2% overlap region was included for stitching, resulting in datasets with a typical logical size of (X, Y, Z)∼(4000, 7500, 100) pixels. In pre-processing, confocal data was cropped to exactly the area of the microelectrode array and downsampled by a factor of 2 in X,Y. The confocal datasets thus had a final logical size before analysis of (X, Y, Z)=(1792, 3584, 100), with pixel sizes of (X, Y, Z)=(1.428, 1.428, 1.514) *µ*m/px. The logical size of the (X,Y) dimensions in the aligned confocal datasets is consequently 7x the dimensions of the microelectrode array datasets, which have shape (X, Y) = (256, 512).

Additional information for the imaging settings unique to each species/application are featured in Table S2.

## ACKNOWLEDGMENTS

This work was supported by the National Science Foundation under Grant No. 2027108. J.W.L. acknowledges support from NIGMS 1R35GM142584-01 and the Burroughs Wellcome Fund. J.K.R. acknowledges the Hazeltine Innovation Award from the Brown University School of Engineering.

J.T.I., J.W.L., and J.K.R. conceived of the research; J.T.I. performed experiments; J.T.I. analyzed data; K.H. and J.K.R. designed the CMOS chip; J.T.I. made figures; all authors wrote and edited the manuscript.

We acknowledge helpful comments from Munehiro Asally and Letícia Galera-Laporta.

The authors declare no conflict of interest.

