## Supplementary Information for "Non-Optical, Label-free Electrical Capacitance Imaging of Microorganisms"

### SUPPLEMENTARY MATERIAL

**Supplementary Table S1.** Species and Strain Information

| Figures | Strain | Description |
|---|---|---|
| Fig 1<br>Fig 3<br>Fig S3 | JMJ1061 | B. *subtilis* NCIB3610 pBS32(*comI* Q12L) *lacA*::Ppen-mApple-kan<br>*amyE*::PcapB-YFP-spec Δ*sinR*::mls |
| Fig 2(i) | PS216 | B. *subtilis* PS216<br>An undomesticated Slovenian soil isolate [**Durett2013**]. |
| Fig 2(ii) | ATCC14580 | B. *licheniformis* ATCC14580<br>Species type strain [**Jespersen2012**]. |
| Fig 2(iii)<br>Fig 4 | YHK038 | S. *cerevisiae* BY4742 (MAT$\alpha$ his3Δ1, leu2Δ0, met15Δ0, ura3Δ0),<br>with FLO and mating genes knocked out<br>(ΔFLO1, ΔFLO5, ΔFLO9, ΔFLO10, ΔFLO11, ΔSAG1, ΔAGA1, ΔAGA2, ΔFIG2).<br>FLO11 knocked into the URA3 locus for constitutive mNeonGreen expression from the pTEF1 promoter. pTEF1-FLO11-T2a-mNeonGreen-tADH1- |
| Fig 4 | JMJ1222 | B. *subtilis* PS216 *lacA*::Pveg-mScarlet-kan. |
| Fig 5<br>Fig S8 | NCIB3610 | B. *subtilis* NCIB3610<br>Species type strain [**Lilge2021**]. |
| Fig S4 | ATCC13880 | S. *marcescens* ATCC13880<br>Species type strain. |

**Supplementary Table S2.** Sample information relevant to ECI/FL imaging

| Figures | Media | Dye Name | Excitation/ Emission |
|---|---|---|---|
| Fig 1<br>Fig 3 | 1% Agarose MSgg | mApple | 555 nm /(560nm - 839nm) |
| Fig 2(i) | 1% Agarose MSgg | Thioflavin T | 405nm /(420nm - 714nm) |
| Fig 2(ii) | 1% Agarose LB | Thioflavin T | 405nm /(420nm - 714nm) |
| Fig 2(iii) | 1% Agarose YPD | mNeonGreen | 492 nm /(497nm - 748nm) |
| Fig 4 | 1% Agarose YPD | mNeonGreen | 504 nm /(509nm - 574nm) |
| Fig 4 | 1% Agarose YPD | mCherry | 587 nm /(592nm - 750nm) |

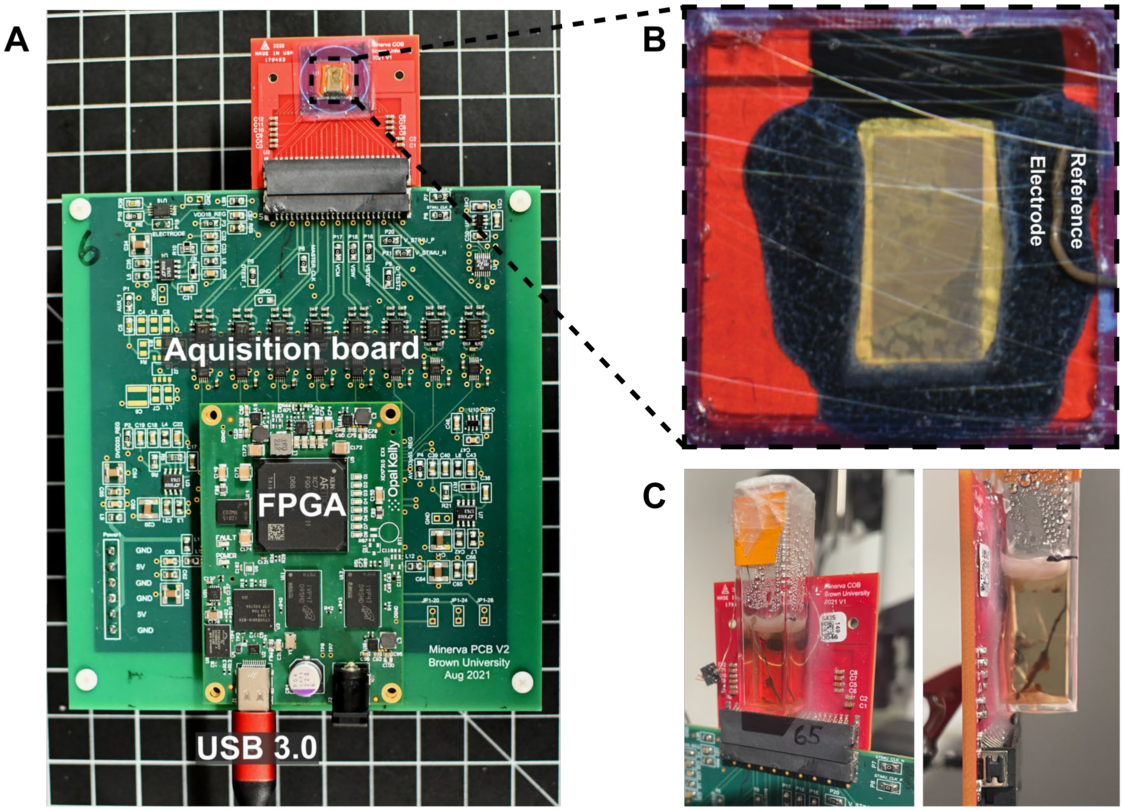


**Supplementary Figure S1. ECI acquisition hardware, and sample preparation** (A) Data from the 131072 CMOS pixels is multiplexed into 8 readout channels, and streamed to a dedicated FPGA via a data-acquisition board. There it is decoded and transmitted via USB connection to a measurement PC, and saved in HDF5 file format. (B) The agarose sample of Fig 1D/E is seen mounted over the CMOS array, with an optional AgCl reference electrode inserted. (C) Views of the cuvette setup used in pellicle culturing, featuring the pellicle of Fig. S8 after several weeks of growth.

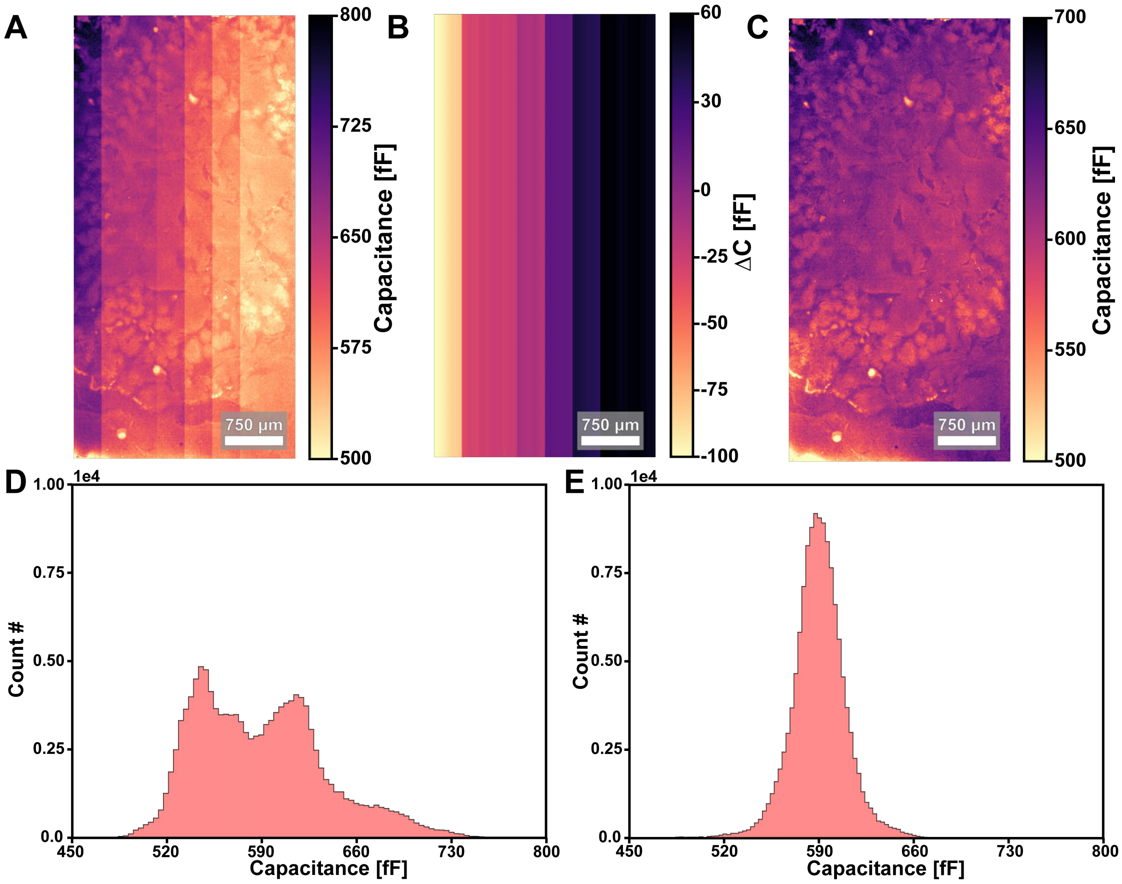


**Supplementary Figure S2. Adjusting for channel readout artefacts** (A) A capacitance image with an offset artefact from the 8 channel CMOS readout. (B) The capacitance correction to be applied to the image and (C) the corrected image. (D) Histogram of capacitance values before (E) and after the correction.

The 8-channel readout introduces slight offset artefacts in the resulting ECI image, that are adjusted for before analysis. This is the only modification to ECI data in this paper, aside from one notable exception in the presentation of data in Fig. 2C (i,ii,iii). Here the capacitance images were anti-aliased through up-sampling by a factor of 16 with bi-linear interpolation (image size was increased from [100,100] to [400,400]).

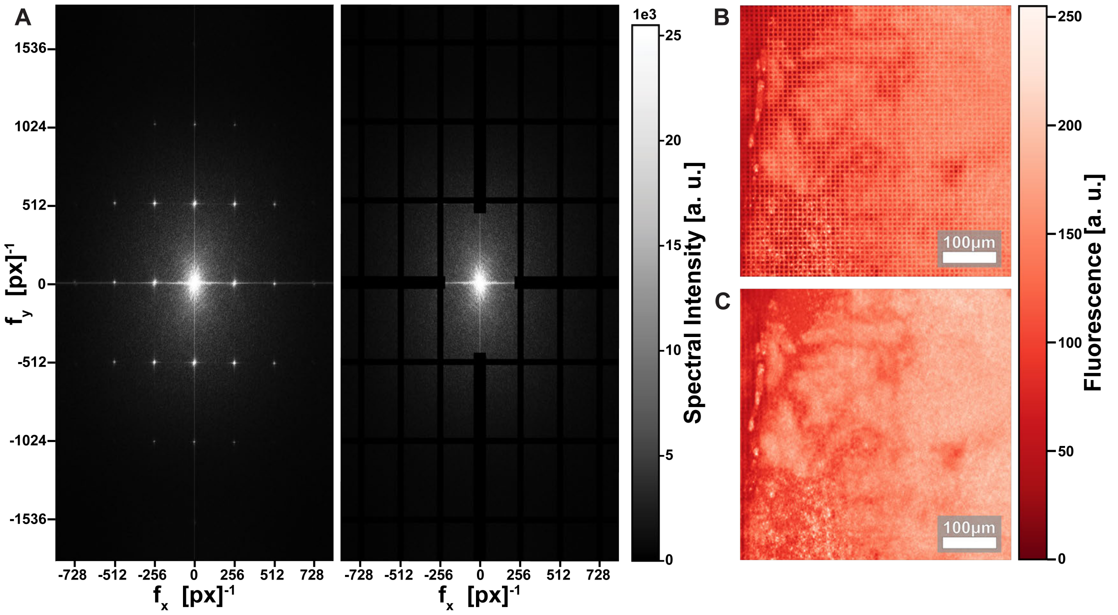


**Supplementary Figure S3. FFT suppression of CMOS features in optical images** (A) The 2D FFT of a confocal image (left) with periodic features showing up at increments determined by the shape of the CMOS array (256,512). (B) A simple grid filter is applied in the frequency domain to suppress CMOS features.(C) The confocal image before (D) and after applying the FFT filter.

By intentionally keeping the wavelength separation between excitation and emission narrow (Table S2), we capture the scattering of light off of the CMOS electrodes, making them visible in the confocal z-stack datasets. The visibility of the electrodes allows the plane of the CMOS surface to be established within the confocal z-stack. This enables the precise (X,Y) alignment and comparison of ECI and confocal datasets, as well as the characterization of biofilm thickness used in Fig 3. However in high resolution fluorescence images of the full CMOS area, the interaction between the stitching of a (8x15) tile grid and the rectangular CMOS pixels visible in the image produces a jarring Moiré effect when viewed on a computer monitor. Thus for the presentation of data in Fig. 1E, the periodic features of the CMOS array were suppressed by applying a simple filter to the image data in the frequency domain. Fig S3 demonstrates the method.

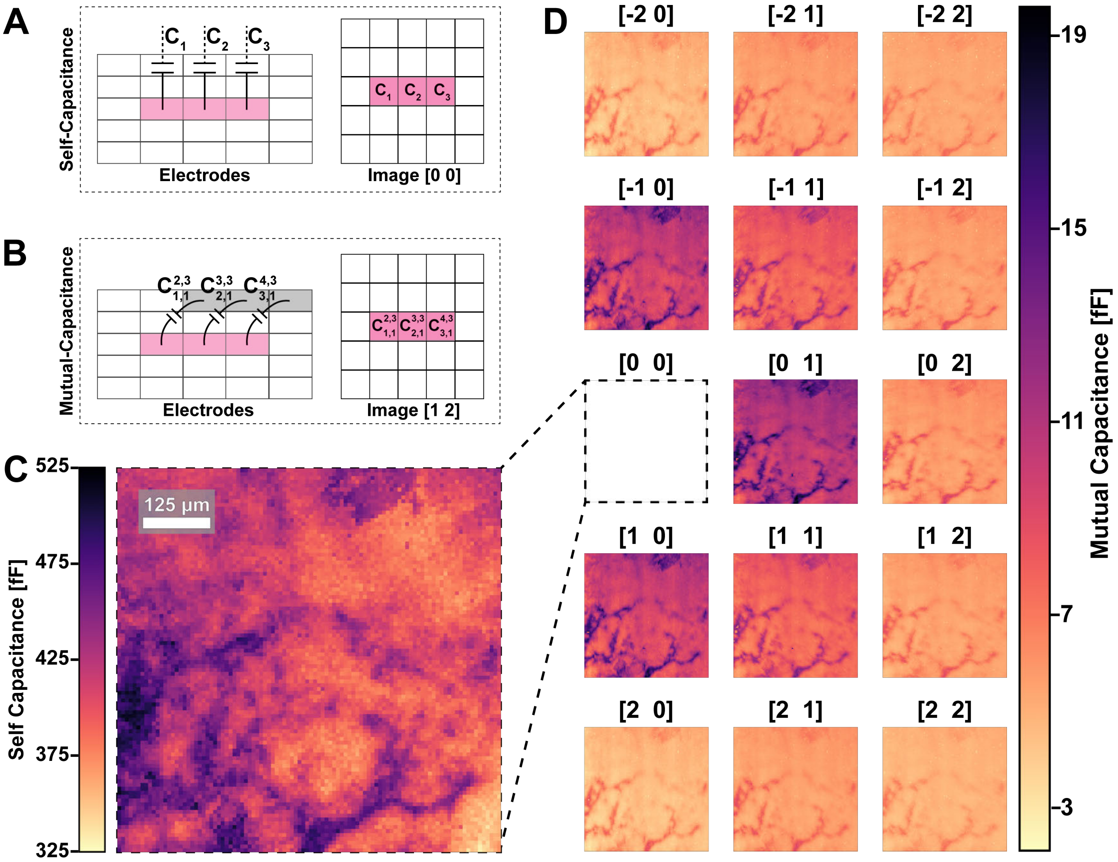


**Supplementary Figure S4. Self vs mutual capacitance modalities** (A) Self-capacitance and (B) mutual capacitance measurements between pixels are methods to take ECI images without a reference electrode. (C) Self-capacitance image of a *Serratia marcescens* colony. (D) Mutual capacitance images of the same colony using the capacitance measured between pixel pairs of varied spacing.

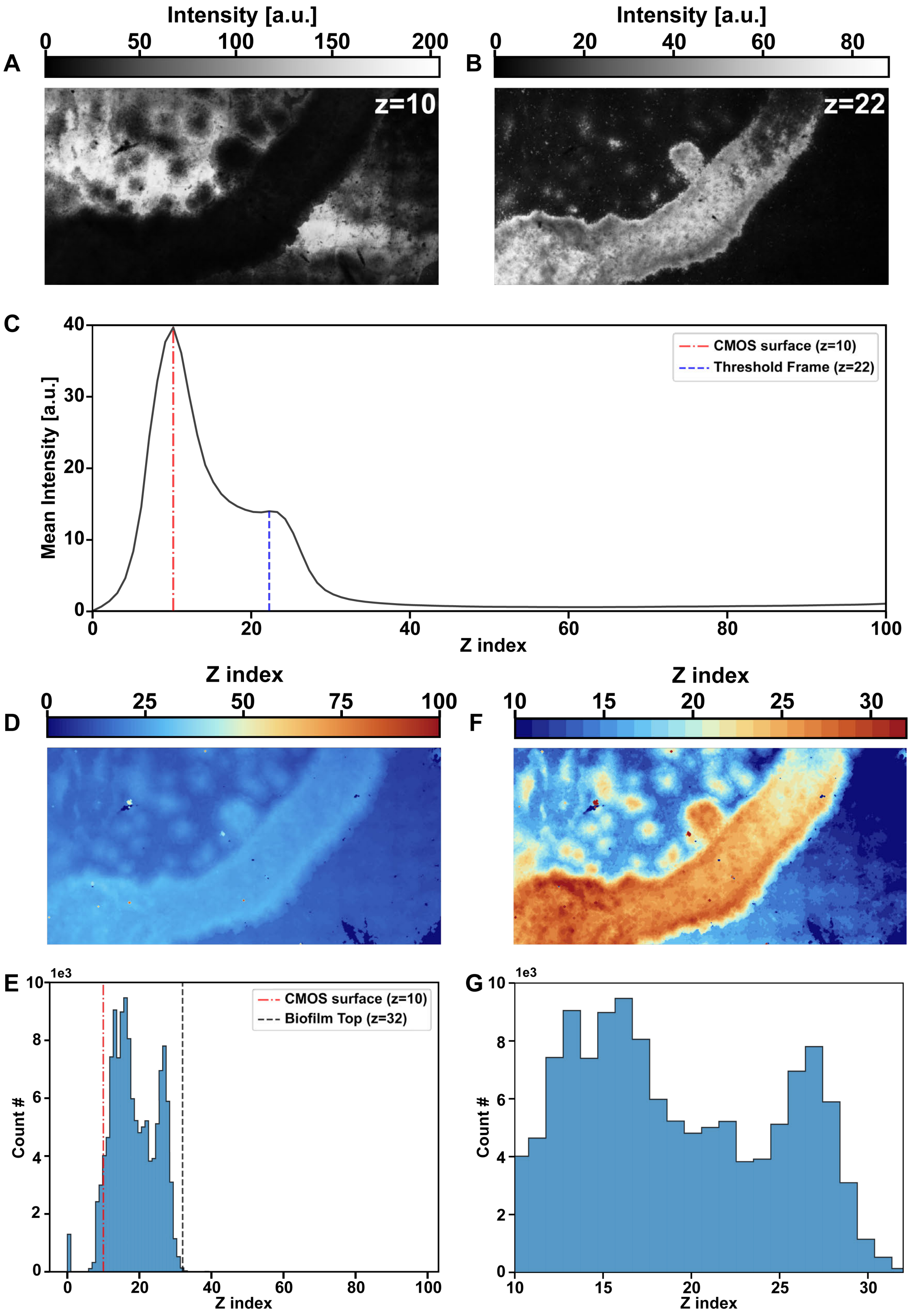


**Supplementary Figure S5. Mapping biofilm thickness with 3D segmentation** (A) The plane of the CMOS array and (B) a frame for thresholding are determined from the mean intensity profile (C) across z indices. (D) A 2D heatmap of the maximum z positions where signal is present is produced using the threshold determined in (B). (E) The distribution of z value maxima reveals the highest point of the biofilm, as well as signal outliers detected both beneath the CMOS and above the sample in the media substrate. (F) The final heatmap of biofilm thickness and the corresponding distribution (G) after outliers are suppressed.

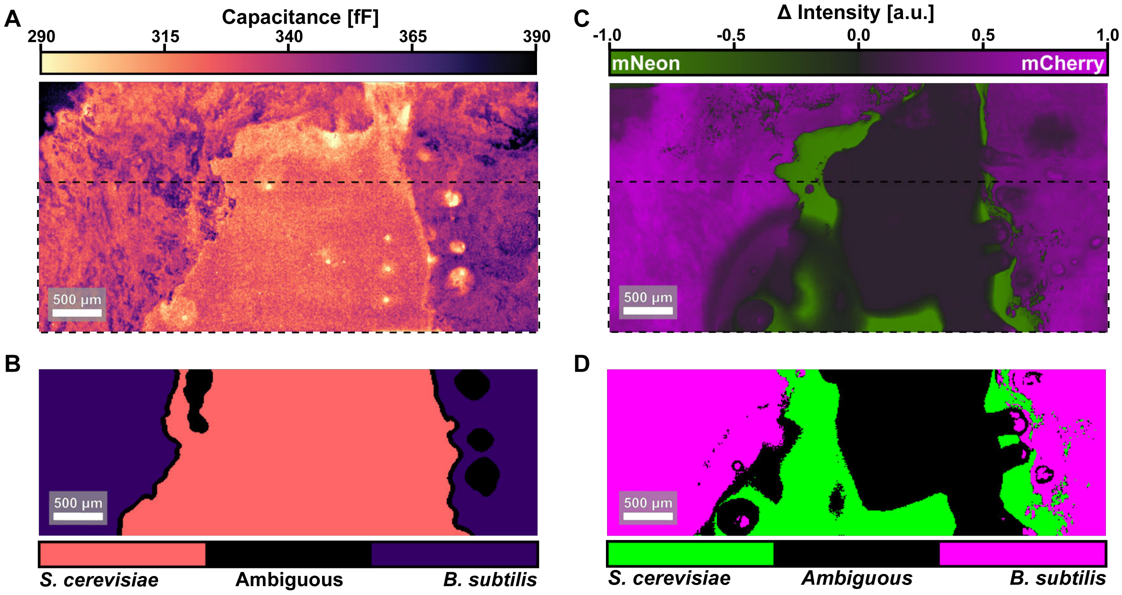


**Supplementary Figure S6. Pattern Segmentation of multi-species ECI data.** ECI data (A) is cropped to a region without media pockets, where we expect to measure only biomass from one of the two species. (B) A mask of the species locations is created manually using the apparent boundaries between the two populations. (C) A multichannel fluorescence image of the same region is used to identify the species of both populations (D). Notably, mNeon expression is weak in the center of the sample, yet the presence biomass there is clear in ECI.

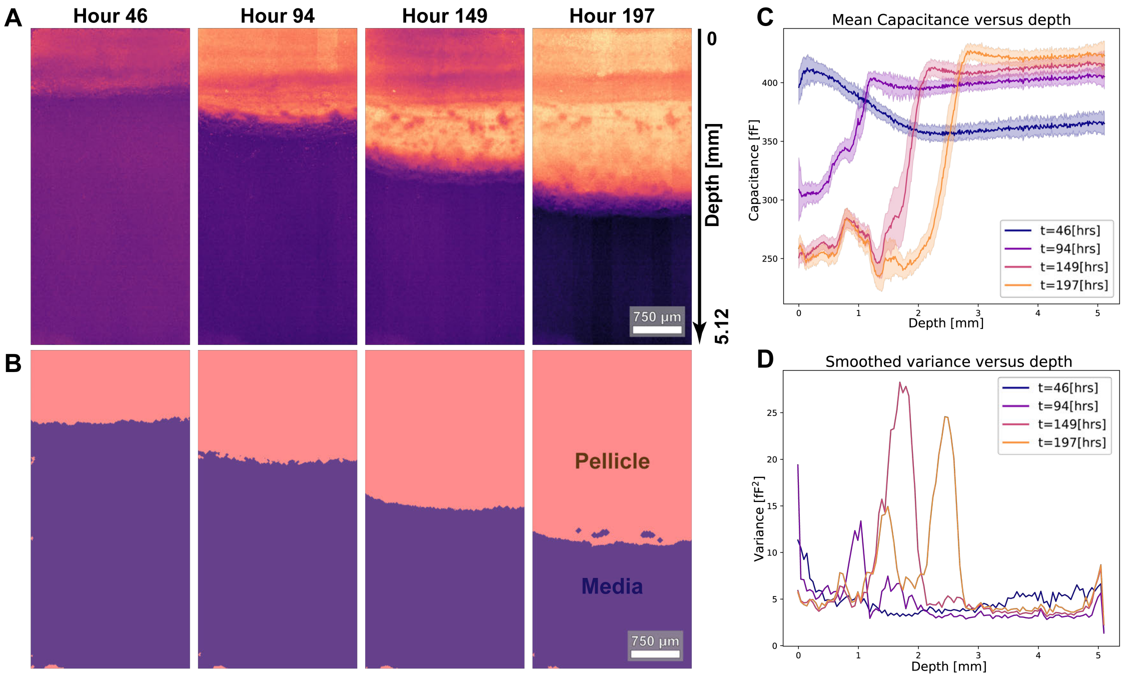


**Supplementary Figure S7.** **Tracking pellicle formation in ECI time-lapse data** (A) ECI time-lapse data from Fig. 5 with the corresponding masks (B) identifying the expanding pellicle region. Traces of the capacitance mean (C) and variance (D) in the ECI images as a function of depth.

#### Long term ECI time-lapse

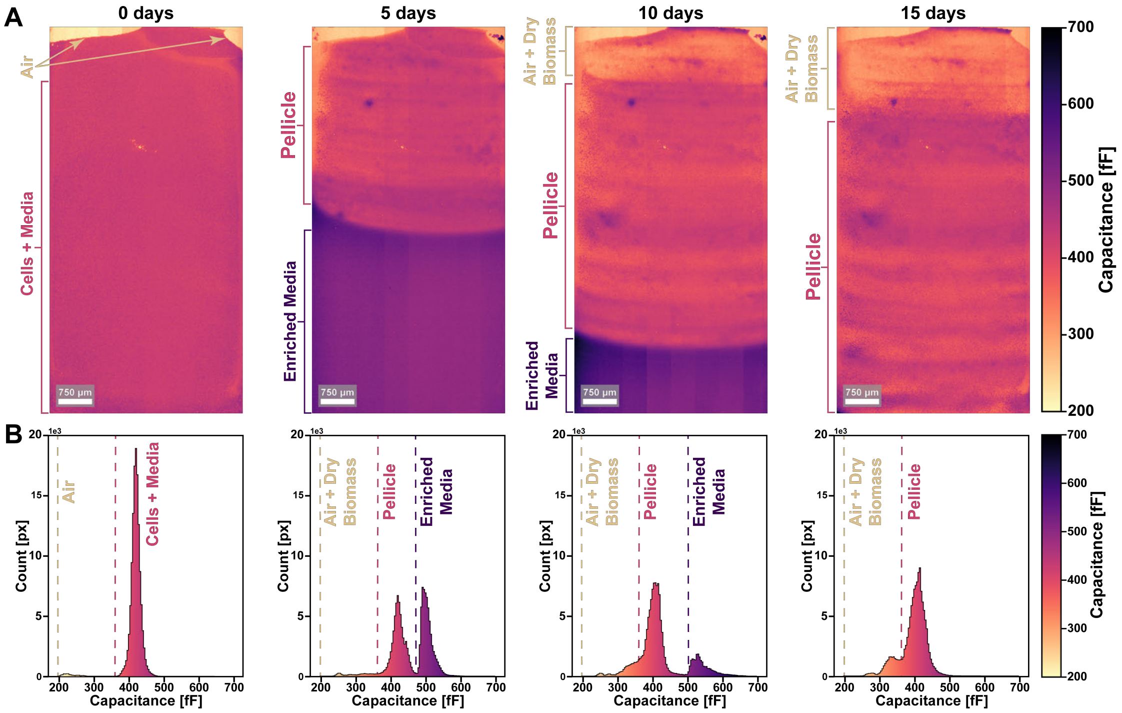


**Supplementary Figure S8. Long term monitoring of pellicle development**(A) ECI data from a 16 day time-lapse captures the dynamics of pellicle formation at the liquid-air interface. (B) Histograms of the capacitance distribution corresponding to each time-point.

In Fig. S8A, we see the initially well mixed cell suspension (day 0) separates to form a pellicle at the air-liquid interface (day 5). Meanwhile, cell activity gradually causes electrolytic enrichment of the surrounding media, due to the continuous excretion of metabolites. As a result, culture media becomes increasingly distinguishable from pellicle biomass in the capacitance distribution (day 5 - 10), with media signal increasing in value relative to the signal of air and biomass (S8B), before eventually disappearing as pellicle growth exceeds the CMOS field of view (day 15). Evaporative losses slowly cause the liquid-air interface in the cuvette to recede downwards (day 10 - 15), which in turn causes biomass at the top of the pellicle to dry against the CMOS sensor. This dry biomass population can be seen across the histograms of Fig. S8B, emerging as a distinct peak between the values associated with the air and pellicle regions.
